# Approaches for integrating heterogeneous RNA-seq data reveals cross-talk between microbes and genes in asthmatic patients

**DOI:** 10.1101/765297

**Authors:** Daniel Spakowicz, Shaoke Lou, Brian Barron, Tianxiao Li, Jose L Gomez, Qing Liu, Nicole Grant, Xiting Yan, George Weinstock, Geoffrey L Chupp, Mark Gerstein

## Abstract

Sputum induction is a non-invasive method to evaluate the airway environment, particularly for asthma. RNA sequencing (RNAseq) can be used on sputum, but it can be challenging to interpret because sputum contains a complex and heterogeneous mixture of human cells and exogenous (microbial) material. In this study, we developed a methodology that integrates dimensionality reduction and statistical modeling to grapple with the heterogeneity. We use this to relate bulk RNAseq data from 115 asthmatic patients with clinical information, microscope images, and single-cell profiles. First, we mapped sputum RNAseq to human and exogenous sources. Next, we decomposed the human reads into cell-expression signatures and fractions of these in each sample; we validated the decomposition using targeted single-cell RNAseq and microscopy. We observed enrichment of immune-system cells (neutrophils, eosinophils, and mast cells) in severe asthmatics. Second, we inferred microbial abundances from the exogenous reads and then associated these with clinical variables -- e.g., *Haemophilus* was associated with increased white blood cell count and *Candida,* with worse lung function. Third, we applied a generative model, Latent Dirichlet allocation (LDA), to identify patterns of gene expression and microbial abundances and relate them to clinical data. Based on this, we developed a method called LDA-link that connects microbes to genes using reduced-dimensionality LDA topics. We found a number of known connections, e.g. between *Haemophilus* and the gene IL1B, which is highly expressed by mast cells. In addition, we identified novel connections, including *Candida* and the calcium-signaling gene CACNA1E, which is highly expressed by eosinophils. These results speak to the mechanism by which gene-microbe interactions contribute to asthma and define a strategy for making inferences in heterogeneous and noisy RNAseq datasets.

## INTRODUCTION

### Linking high-dimensional, heterogeneous datasets

RNA sequencing (RNAseq) has become a standard method of analyzing complex communities. Depending on the sample type, these data can be very heterogeneous. A key problem tackled in this paper is dealing with the heterogeneity and noise in RNAseq data in complex samples such as sputum. This can be appreciated by comparing sputum RNAseq to a more traditional experiment, e.g. blood RNAseq, where the sample can be collected consistently and that contains relatively well-defined cell types (Figure 1). In blood, the vast majority of RNAseq reads align to the human genome, and the goal is often to relate the expression of the genes to a phenotype. By contrast, sputum may be less consistently collected, its cell types are less defined, and it may contain RNA from microbes and other organisms that act as cryptic indicators of the environment. This combination of variables and dimensions often requires researchers to collapse the dimensions to appropriately de-noise the analysis. Here, we present such a strategy that uses a number of supervised and unsupervised techniques such as single-cell signatures and latent Dirichlet allocation (LDA). These techniques can produce a low-dimensional representation of common groups of genes, microbes, or other features that tend to increase or decrease in abundance together. Our approach is useful when the heterogeneity comes from the sample type (e.g., sputum) and especially when the samples derive from a heterogeneous population of individuals, such as patients with asthma.

**Figure 1.**
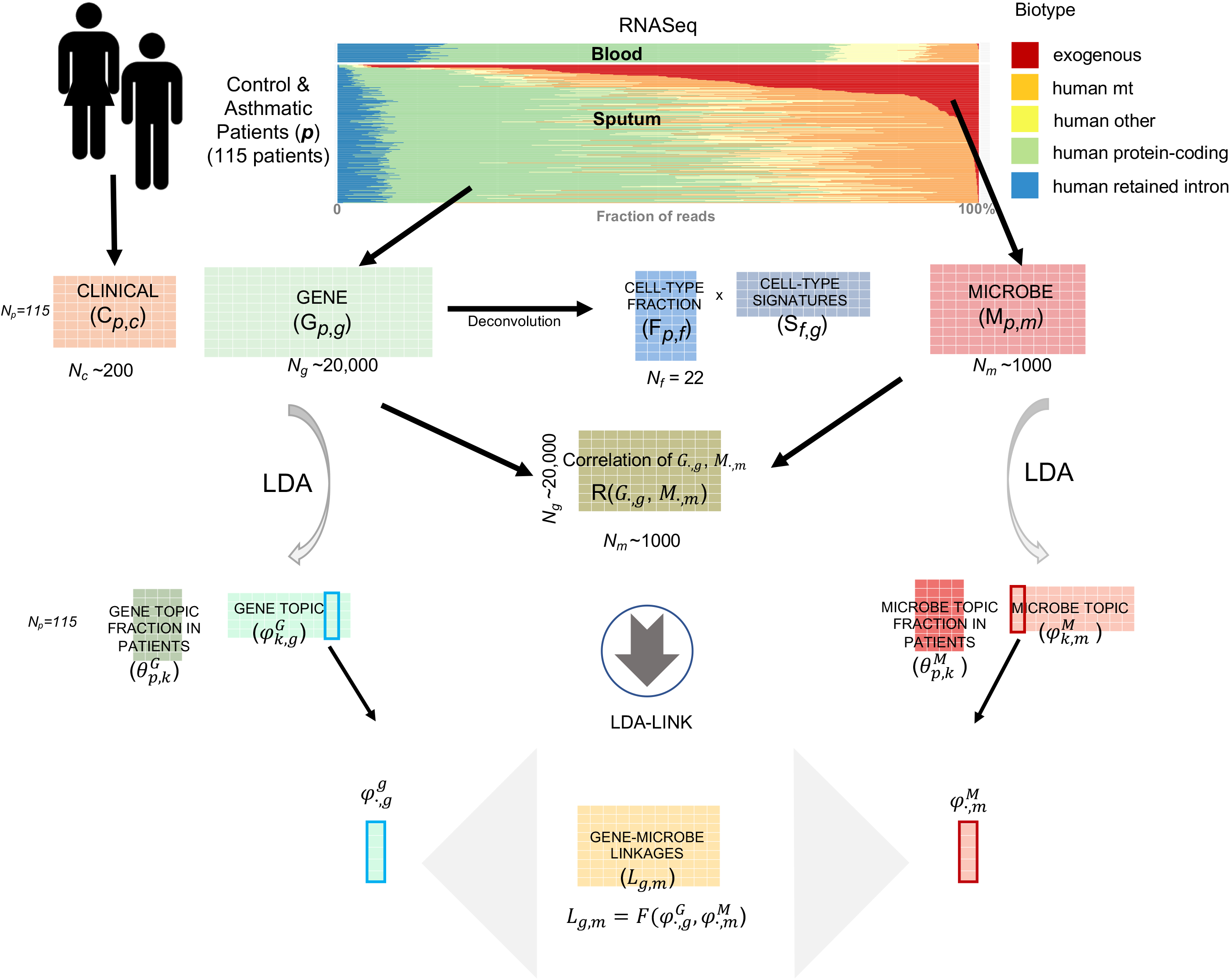
RNAseq alignment summary for control and asthmatic sputum, showing fractions of reads that aligned to different biotypes. Alignments to the protein-coding biotype were used to generate the gene expression matrix (G), which was then deconvolved into a cell fraction matrix (F) and cell expression (E). The exogenous reads were used to generate a microbial profile matrix (M). These matrices were then related to the clinical phenotype matrix (P) for biological insight.

### Interactions between the host and microbes in the lung

Asthma is a disease of the airway that can present with many clinical phenotypes. Much work has focused on identifying subgroups of the disease and how each subgroup responds to treatment. For example, Yan et al. introduced transcriptional endotypes of asthma and the Severe Asthma Respiratory Phenotype consortium defined five subtypes of asthma [1]. Some of these subgroups respond differently to environmental and microbial triggers, such as fungal spores. Some fungi have well-defined effects in asthma, but the role of many microbes remains contentious. A simplified model assigns microbes to one of three categories: pathogenic organisms that cause inflammation, beneficial organisms that reduce inflammation, and those that have no effect on inflammation. The majority of the organisms in the lungs are expected to have no effect, and severe asthmatics are expected to have more pathogenic and fewer beneficial microbes.

### Inferring immune cell fractions from RNAseq data

The pathology of microbes is often inferred by the number and type of immune cells observed in samples, such as sputum total leukocyte counts [2, 3]. A standard method for counting immune cells in sputum samples uses microscopy, but the resolution is limited to a few cell types [4]. Other cell-counting methods such as flow-sorting can be challenging because of the viscosity and highly variable cell numbers in sputum. An alternative strategy uses cell-type specific expression patterns to deconvolve RNAseq reads from mixtures of cells into fractions of different immune cells [5]. This deconvolution also effectively de-noises heterogeneous datasets by greatly reducing the number of dimensions. Importantly, the RNA needed for this analysis can be purified without poly-A enrichment– here, we use human ribosomal RNA knockdown – which allows for the simultaneous analysis of microbial and human transcripts.

### Supervised deconvolution and the microbiome

While deconvolution to cell fractions effectively de-noises human RNAseq data, an equivalent method does not exist for microbes. Although we can map microbe reads onto their genomes, this approach is imperfect because the genome databases are incomplete and assigning a read to a single genome can be complicated if it matches more than one equally well. One can reduce the dimensions by collapsing microbial strains to different taxonomic ranks (e.g., genus or family); however, taxonomy is notoriously imprecise at defining behavior. For example, many bacteria in the genus *Escherichia* are human commensals, whereas *Escherichia coli* OH157:H7 causes hemorrhagic colitis. Alternatively, one can group sequences by the metabolic pathways observed, although this requires high-depth sequencing. Here, we propose a method to reduce the dimensionality of microbes by first linking the microbes to human genes, and then applying the relatively well-defined gene dimensionality-reduction methods (e.g., deconvolution to cell types).

In this paper, we use RNAseq of sputum samples from asthmatic patients to demonstrate dimensionality-reduction strategies and identify microbe-host relationships. We map RNAseq reads onto human or microbial genomes and relate the resulting abundance matrices to each other and to clinical data. Further, we deconvolve the human reads into fractions of the various cell types that make up sputum. Finally, we relate the human genes and microbes using a method we call LDA-link, which identifies relationships between genes, microbes, and cell types. These methods represent a general strategy for dealing with heterogeneous RNAseq data that is applicable to other sample types beyond sputum.

## RESULTS

### Sequencing and processing with the extracellular RNA processing toolkit (exceRpt) pipeline

We collected induced sputum samples from 115 patients with heterogeneous asthma phenotypes and sequenced these sample using RNAseq. The median read depth per sample was 47.5 million, which meets depth recommendations for analyses of this type [6]. We processed these reads through the exceRpt pipeline [7], which conservatively matches reads to genomes in a sequential order designed to reduce experimental artifacts. In brief, we first aligned the quality filtered reads to the UniVec database of common laboratory contaminants^2^, and then aligned the remaining reads to human ribosomal sequences before aligning them to the human genome. We excluded samples with a low ratio of transcript alignments to intergenic sequence alignments, and then aligned the remaining reads to the comparably large sequence space of non-human genomes. We first aligned reads to the relatively well-curated ribosomal databases of bacteria, fungi, and archaea (e.g., Ribosomal Database Project^3^) and then to curated genomes of bacteria, fungi, viruses, plants, and animals. The percent of reads mapping to different biotypes was highly heterogeneous; a median of 60% of the reads aligned to the human reference genome and 50% to annotated transcripts (Figure 1, green bars). A median of 0.7% of the input reads aligned to exogenous sources, with some samples containing as much as 28.1% exogenous reads. As a control, we applied the same protocol to blood samples, which demonstrated more homogeneity than sputum (Figure 1, top, “blood”).

### Overview of the analysis approach

The goal of the analysis was to infer meaningful relationships between the numbers and origins of the RNAseq reads and relate them to clinical phenotypes. We conceptualized the clinical information and RNAseq alignments as a series of tables (**Figure 1**). The clinical table includes patient data collected at the clinic, ***C***, including age, weight, lung function tests, etc, with rows indexed by patient (*p*) and roughly 200 clinical variables (*N_c_*). Alignments to human protein-coding regions created the gene table, ***G***, with *N_p_* rows, as above, and roughly 20,000 genes (*N_g_*). Alignments to exogenous genomes created the microbe table (***M***) with *N_p_* rows and roughly 1,000 microbes (*N_m_*). Given these three tables (***C, G***, and ***M***), the basic analysis framework is to correlate columns or rows within or between tables. We represent this by a matrix of correlations, **R**(***X·,_i,_, Y·_j_***), where ***X·,_i_*** is the *i*^th^ column of table ***X*** and ***X·,_j_*** is the *j*^th^ column of table *Y*. This correlation is summed over the other index, usually p. For example, we test the relationship between age and the abundance of each microbe **R**(***C·,_age_, M·,_m_***) across all patients. Similarly, we correlate the expression of a gene (e.g., *TLR4*) with microbe *Candida* **R**(***G.,_TLR4_, M·,_Candida_***).

Individual correlations can be difficult to interpret, particularly in heterogeneous, sparse, or noisy datasets. Organizing the genes into relevant pathways or cell types can reduce the dimensionality and de-noise the analysis. To this end, we deconvolved ***G*** (***N_p_ × N_g_***) into a cell-type fraction table, ***F*** (***N_p_ × N_f_***), and a cell-type signatures table, ***S*** (***N_f_ × N_g_***). However, an analogous supervised method does not exist for the microbes. Therefore, we applied an unsupervised dimensionality-reduction approach, latent dirichlet allocation (LDA), which provides a topic distributions in patients (***θ***^*G*^, ***N_p_× N_k_***) across a smaller number (***N_k_***=10) of topics and gene topic (and ***φ**^G^*, ***N_k_ × N_g_***). This can also be done to the microbe table M and get ***θ**^M^* and ***φ**^M^*, and the gene and microbe topic can be correlated (e.g. 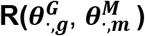 over all patients).

The framework described above is useful for identifying linear relationships, but non-linear relationships are also possible. For example, a microbe sensed by a human immune cell could lead to the activation of a transcription factor and the expression of several genes, each of which would have a non-linear relationship to microbe abundance. To identify such relationships, we applied a non-linear ensemble learning algorithm [8, 9], using the de-noised inputs for each gene and microbe (***φ**^G^* and ***φ**^M^*). We call this method LDA-link. Further, we relate the gene and microbe links identified to cell fractions and thereby relate how the host is responding to microbes with regards to immune cell type response with a particular gene.

### Analysis of human-aligned reads

Working toward the hypothesis that we can conceptualize human-aligned sputum RNAseq reads as a mixture of immune cell types, each with a distinct expression profile, we deconvolved the Gene table (***G***) into a table of fractions of component cells type (***F***) and cognate cell-type signatures (***S***) by solving the formula ***G ~ F * S***. This method relies on knowing the signature gene-set in each cell type, which derived from the blood immune cell high quality profiles. To validate that we could apply these cell expression profiles to sputum, we generated several additional datasets including single-cell RNAseq (scRNAseq), microscopy, and unsupervised decomposition, and then compared the results to the deconvolution table ***F***. (Figure 2A, schema).

**Figure 2.**
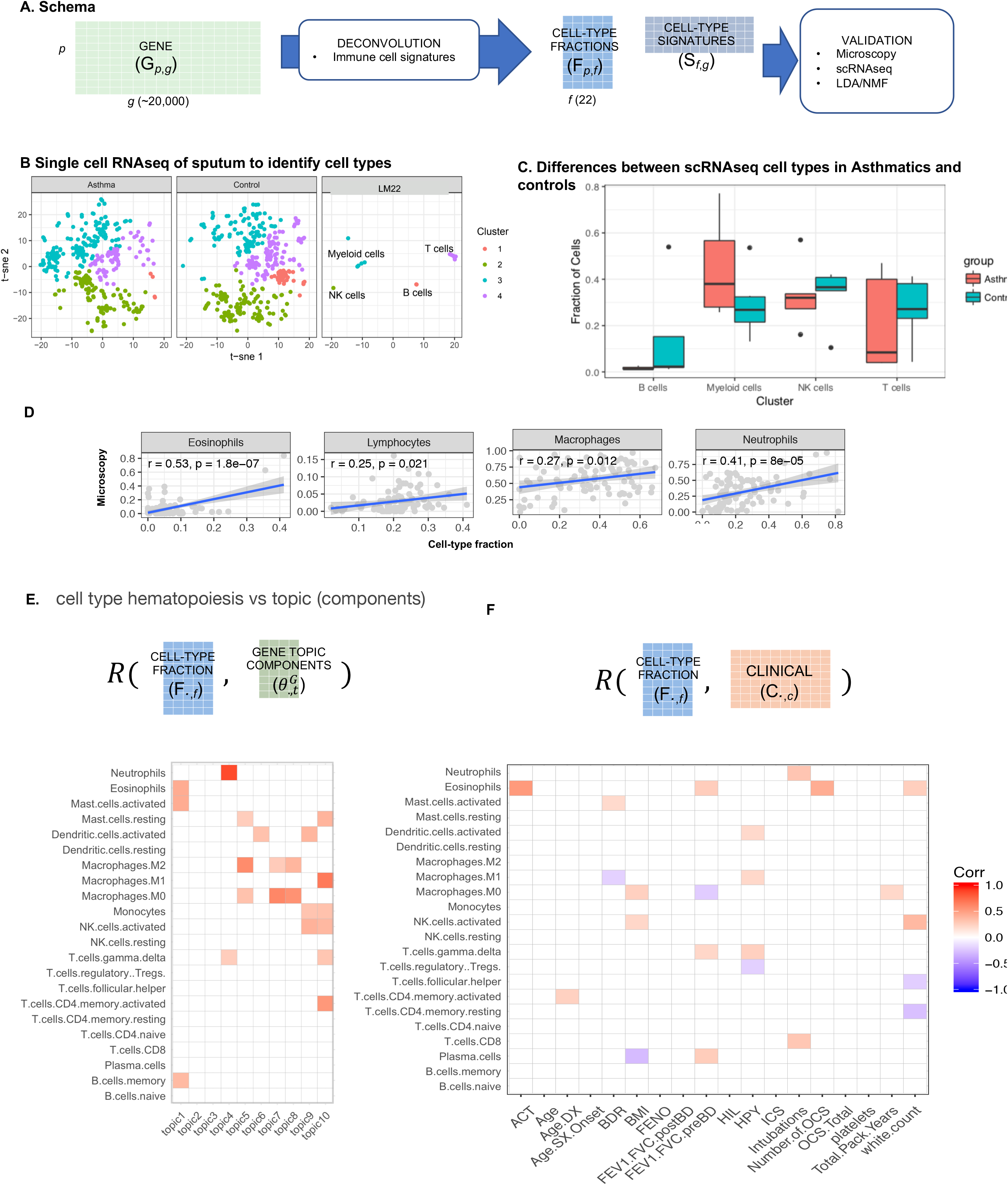
Deconvolution of RNAseq human reads into cell fractions using cell signature deconvolution. A) Schematic showing the imputation of a cell fraction matrix and cell-specific expression matrix. B) Imputed cell fractions were validated using microscopy; Cell fractions were then correlated with SARP cluster for two major cell type: (C) Machrophases.M0 and (D) Mast cell activiated. E) the cell fraction of LM22 gene signature deconvolution are correlated with the topic distribution of samples from LDA analysis. F) G) tSNE analysis and clustering using single cell RNAseq from Asthmatic patient and control. H) The fraction of single cells for different cell types clusters between Asthmatic patient and Controls.

### Evaluation of deconvolution results by scRNAseq

First, we performed scRNAseq on a cohort of similar sputum samples (five control and five asthmatic patients). The single-cell sequences clustered into four groups (Figure 2B, first and second panels). To determine whether the reference profiles that we used to deconvolve the bulk RNAseq recapitulate those found in the single-cell clusters, we co-clustered the reference profiles with the scRNAseq data (Figure 2B, third panel). The reference profiles split into the groups by lineage; for example, those in the lymphoid progenitor line coclustered with cluster 2, and the myeloblast progenitor line co-clustered with cluster 4. This result suggests that the reference profiles accurately represent the cell types in sputum. The myeloid lineage cluster showed a significant difference in the number of cells between asthmatics and controls (Figure 2C). From this analysis, we concluded that (1) the blood-derived cell profiles appropriately fit the sputum cell types and (2) no additional cell types are needed to deconvolve the sputum bulk RNAseq data.

### Evaluation of deconvolution results by microscopy

Second, we evaluated a subset of the samples by microscopy and manually counted the number of neutrophils, eosinophils, lymphocytes, and macrophages. We found good agreement with ***F***, when cell counts could be directly compared, i.e. neutrophils and eosinophils were both present in ***F*** and counted by microscopy. In cases where the deconvolution method gave higher resolution, (e.g., M0, M1, and M2 macrophages versus one type of macrophage by microscopy), the aggregation of the relevant columns in *Ff* correlated well with the microscopy counts (Figure 2D).

### Association of cell fractions with clinical features

Having validated the deconvolution of sputum samples (table ***F***), we then correlated the cell fractions with clinical features (***R***(***F._f_, C.,_c_***) for all patients). We found that the changes in fractions of several cell types were highly correlated with clinical features (Figure 2E). For example, the fraction of T-regulatory cells negatively correlated with the number of hospitalizations per year, suggesting a beneficial role of these cells in the management of asthma.

### Evaluation of deconvolution results by unsupervised decomposition

We compared the signal captured by cell-type deconvolution to an unsupervised decomposition method: LDA. Using LDA, we factored the gene expression table into ten topics that conceptually represent gene expression programs. This resulted in a gene-topic-fraction-in-patients table, ***θ**^G^* (***N_p_× N_k_***) with *N_k_*=10 topics, as well as corresponding gene-topic table, ***φ*** (***N_k_ × N_g_***), that are analogous to the supervised deconvolution tables ***F*** and ***S***. We correlated the cell-type fractions table with the gene topics fraction table (**R**(***F.,_f_, θ._k_***) for all patients, and found agreement between LDA and the cell-signature-based deconvolution for only the most prominent cell type, neutrophils (Figure 2D, topic 4). The top genes associated with topic 4 were enriched in the neutrophil chemotaxis pathway (Figure S8 B).

However, the remaining topics were comprised of multiple cell types. This suggests that LDA can identify distinct but partially overlapping features in ***G***. According to the clustering of ***θ**^G^*, a subgroup of severely asthmatic patients was highly correlated with topic four (**Figure S8A**). The top-weighted genes in topic 4 were enriched for the pathways “neutrophil chemotaxis” and “asthma-related genes” (**Figure S8B**). These pathways were not enriched in the analogous cell-type-signatures table ***S***, suggesting that LDA topics are distinct from the cell-type signatures, but are also clinically relevant. Moreover, the top-weighted genes in topic 1 of the gene topic components table were mitochondrial genes, and topic 1 was strongly correlated with age. This link shows strong support in the literature, as reactive oxygen species produced by the mitochondria reduce their function over time [10]; however, we did not observe this relationship for any cells in the cell-type-fractions table (***F***). Another method using a very different algorithm than LDA, non-negative matrix factorization (NMF), showed strong agreement with LDA (Figure S2, Nmf.1). This supports the use of supervised deconvolution methods as picking out interpretable signals that are different than those identified by unsupervised methods. Unsupervised decomposition should be considered a set of features distinct from those found through deconvolution.

### Analysis of exogenous reads

After filtering out contaminants and human reads, we assembled the set of reads that aligned to exogenous genomes into a Microbe table (***M***). The exogenous sequences aligned to mostly bacteria and fungi, although we also observed a few arthropod and helminth reads (**Supplemental Table X**). The dominant phyla observed were from the bacterial kingdom: Proteobacteria, Firmicutes, and then Bacteroidetes. The abundance of Proteobacteria is in contrast to observations from the gut where Bacterioidetes predominate [11]. Also notable was the presence of two phyla of fungi among the eight most abundant overall, although this was in lower abundance than many of the bacterial phyla.

### Microbes correlations with clinical information and cell fractions

We correlated the microbe abundances to clinical information (***R***(***M·,_m_, C·,_c_***) for all patients) (Figure 3A). *Haemophilus* was associated with increased total white blood cell numbers, as has been described previously [12]. *Candida* was associated with worse lung function test results (e.g., forced expiratory volume and forced vital capacity), which supports the association with a severe form of asthma characterized by eosinophilia [13].

**Figure 3.**
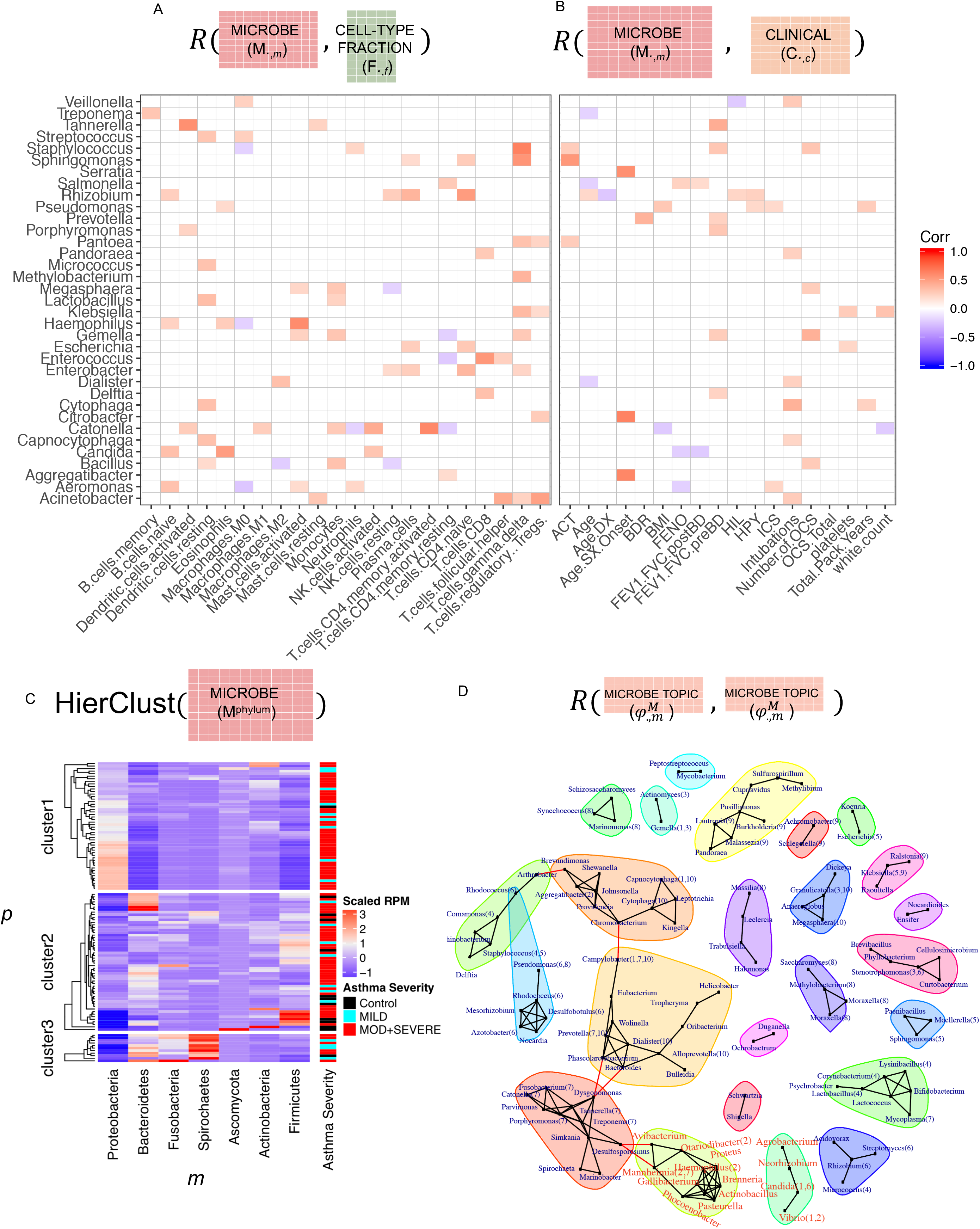
Exogenous RNAseq analysis. (A). The correlations between microbes abundance and cell fraction based on LM22 signature (B) The correlation between microbes abundance and clinical information. (C) The microbes abundance shows clear patterns that associated with Asthmatic severity. (D) The co-abundance network and overlay with the associated topics of microbes.

We next correlated microbe abundances to human immune cell fractions (***R***(***M·,_m_, F·,_f_***) for all patients) (**Figure 3B**). Several correlations demonstrated results with strong literature precedence. For example, studies have previously shown that *Haemophilus* associates with eosinophilia [14], and we observed a significant correlation between *Haemophilus* and the fraction of eosinophils. We also observed a significant correlation between *Haemophilus* and activated mast cells, suggesting an alternative route to *Haemophilus-induced* inflammation [15]. Moreover, the fungal genus *Candida* was also significantly correlated with eosinophils, even more strongly than *Haemophilus.* Pulmonary candidiasis has long been associated with allergic bronchial asthma and inflammation [16], however few lung microbiome studies have examined both bacterial and fungal signals. This highlights the need for a more comprehensive search of the lung microbiome and demonstrates the power of an RNAseq-based method that can report on all kingdoms with the same sample preparation.

### Dimensionality reduction for microbes: clustering and networks

We attempted to de-noise the microbe table (***M^Phylum^***) with a variety of dimensionality-reduction techniques. First, we collapsed the microbes by taxonomy, grouping them to the rank of phylum, and then hierarchically cluster the patients based on their phylum abundance (Figure 3C ***HierClust***(***M^phylum^***)). The hierarchical clustering showed that the phylum distributions formed three clusters of patients. We related these clusters to the clinical variable “asthma severity” and observed that cluster 2 was enriched for patients identified as having moderate or severe asthma. This cluster was characterized by the highest relative abundance of the phylum Proteobacteria (Figure 3C). Notably, the genus Haemophilus belongs to this phylum, consistent with the correlations observed at the genus rank (Figures 3A, 3B).

Similarly, we could de-noise the microbe table using a co-abundance network, by correlating the genus-level abundances (***R***(***M·,_m_, M·,_m_***) and identifying significant modules (**Supplemental Figure Z**). An interpretation of these modules is that they define metabolic niches, where microbes either directly compete for metabolites or there is interdependency in metabolite production. Such networks could be created from other tables, such as the topic distribution of microbes 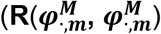 for all the topics) (**Figure 3D**). These modules represent another unit that could be related to the clinical information (**C**) and the cell-type fractions (***F***).

### LDA-link for the identification of links between genes and microbes

How much cross-talk exists between microbes and human cells in the airway remains contentious [17]. We feel this is partly due to the heterogeneous and noisy data from airway samples, where it is often difficult to find strong correlations using standard algorithms. We therefore sought to link genes to microbes via a new method called LDA-link.

LDA-link connects genes to microbes using a combination of linear correlation, unsupervised decomposition and an ensemble learning classifier. We hypothesized that the only strongest gene and microbe correlations would be observable through the noise in the RNAseq data. Therefore, we used these strong links as a training set to find other links, after taking steps to reduce the noise in the data. We reduced the noise using LDA and then identified links using a random forest classifier, described in more detail below and in the methods section.

To define the training, set we first related columns between the gene and microbe tables (**R**(*G.,_a_, M,_m_*)), yielding many low-scoring correlations. However, a relatively small number were strong (**R** > 0.4) and highly significant (p < 1E-5 after FDR correction) (**Figure 4A**). We selected the very strong correlations as true-positive links between genes and microbes in the training set, and non-correlated pairs (−0.05 < **R** < 0.05) as true-negative links. The genes involved in these strong correlations were enriched for pathways related to microbial interactions in the airway, including “Asthma & Bronchial Hypersensitivity” and “Respiratory Syncytial Virus Bronchiolitis” (**Figure 4B**), suggesting that the small set of strong linear correlations were relevant to asthma.

**Figure 4.**
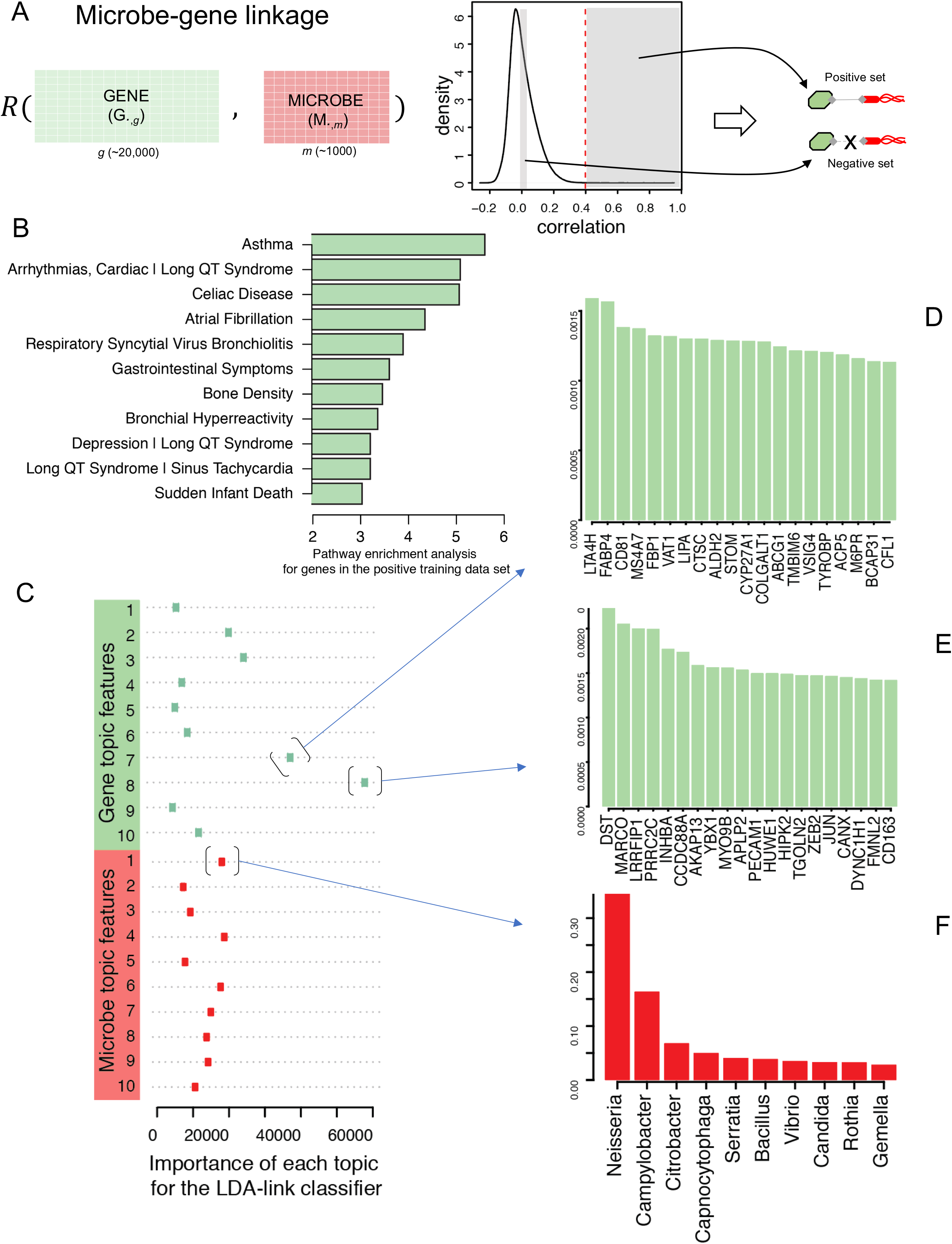
Prediction of cross-talk between microbe and gene. (A) The diagram to combine linear and LDA-based non-linear algorithms to identify gene microbe linkages. (B) simple correlation to identify strong linkages between microbes and genes. Gene set over represent analysis for genes. X-axis is the -log(p-value). (C) the importance of features (LDA topics for gene and microbes) in the RandomForest model by Gini index. The top 20 associated gene in topics 8 (D) and topic 7 (E) of genes, and topic 1 (F) of microbes.

Next, we trained a random forest classifier on the linear correlations described above. To reduce the noise in the data, the features used as inputs to the classifier were the LDA topics for each gene and microbe 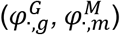. That is, for each gene-microbe pair, we concatenated the gene and microbe topics into a single vector (length 20). The Gini index showed the most important features in defining links between genes and microbes were gene topics #7 and #8, and microbe topic #1 (**Figure 4 C-F**). The genes that comprise the most influential gene topic #8, are enriched for the pathway “Inflammatory Response”, and specifically the cytokines IL2 and IL6. It is tempting to speculate that these genes are strong predictors of a link between genes and microbes because they indicate when the presence of a microbe has triggered an inflammatory response.

### Cross-talk between genes and microbes defined by LDA-link

LDA-link identified connections between genes and microbes reported elsewhere in the literature as well as novel observations. A bipartite graph summarizes a subset of the connections, showing in most cases several genes linked to each microbe (**Figure 5A**, for a complete list see **Supplemental Table X**). Notably, both fungi and bacteria showed these links, further highlighting the need to evaluate more than bacteria when performing microbiome experiments in the airway. The gene lactotransferrin was linked to *Aeromonas,* which has been associated with gastroenteritis and skin infections and has been previously reported to bind lactoferrin [18]. *Burkholderia,* a gram-negative bacterial genus, is recognized as an important pathogen in the mucus-filled lungs of patients with cystic fibrosis; it was linked to gene MUC6, which encodes a secreted protein responsible for the production of mucin [19]. *Haemophilus* was observed to be linked to NFKB Inhibitor Zeta, which is induced by the bacterial cell wall component lipopolysaccharide [20]. In addition, *Haemophilus* was linked to the cytokine interleukin 1 beta (IL1B), an important mediator of the inflammatory response. IL1B hypersensitivity is a hallmark of the asthma phenotype. *Pasteurella* was also linked to IL1B, and its toxin has been shown to induce expression of IL1B [21]. In addition to single gene-microbe pairs, we layered on pathway and cell deconvolution data to identify larger-scale effects of microbes.

**Figure 5.**
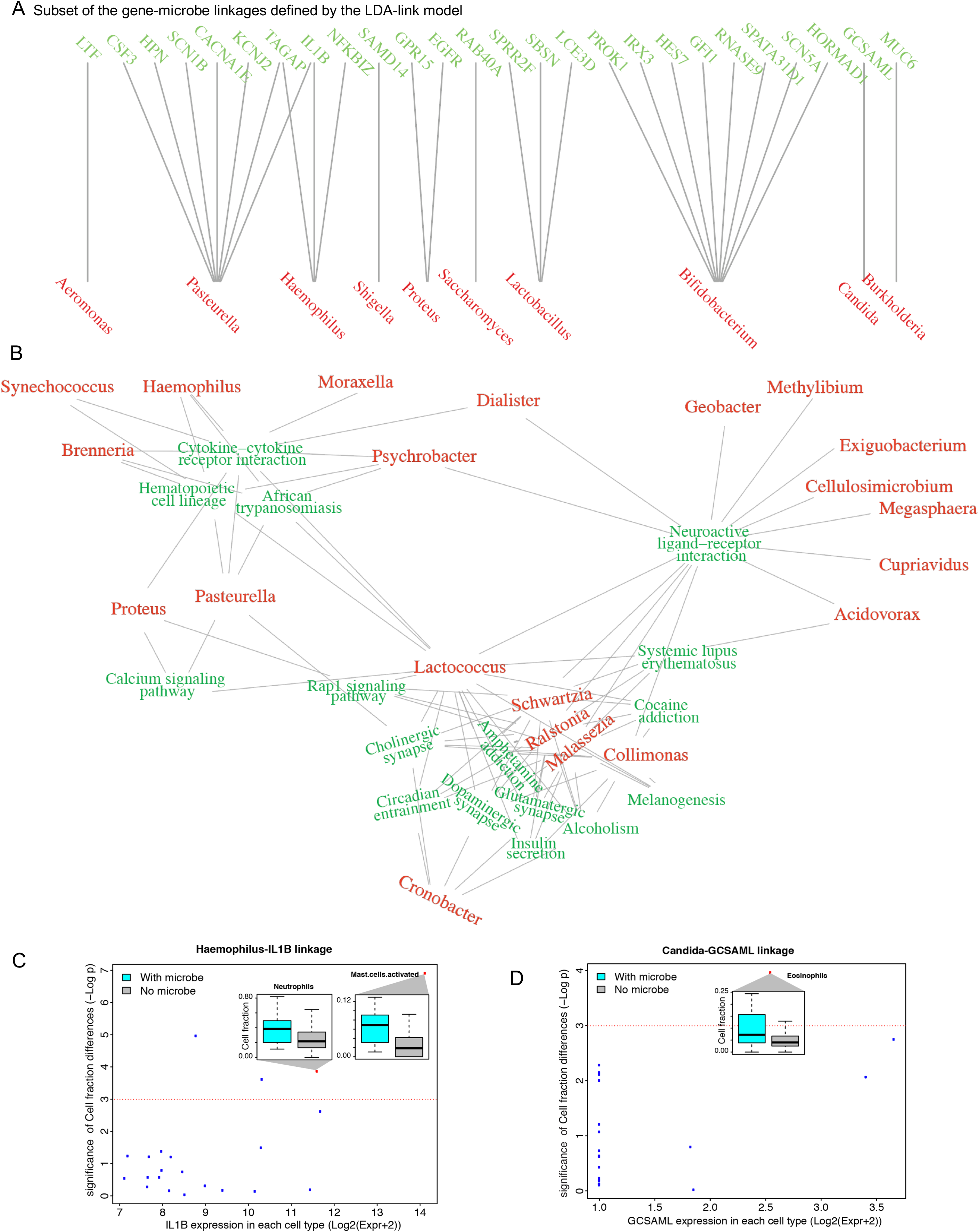
The linkage between microbes and genes reflects the heterogeneity of different cell types. (A). Linkages between microbes and genes. (B) The linkages indicated by the cell proportion of certain types.

Microbes were linked to genes that are enriched in pathways relating to auto-immunity and inflammation as well as cytokine receptors and their interactions (**Figure 5B**). The microbes associated with cytokine pathways included *Synechococcus, Lactococcus, Dialister, Psychrobacter, Moraxella, Brenneria, Proteus, Haemophilus,* and *Pasteurella.* In addition, we related the cell-type signatures table (***S**_f,_g__* to identify the immune cell types that are related to each microbe (**Figure 5C**). We observed the *Haemophilus-IL1B* linkage in monocytes and mast cells. Samples containing *Haemophilus* triggered more activated mast cells according to its cell fraction (**Figure 5C** inset) [22–25]. Similarly, the fungal genus *Candida* was linked to the gene GCSAML, which was highly expressed by eosinophils. The presence of *Candida* was associated with increased numbers of Eosinophils in the airway.

## DISCUSSION

Heterogeneity and noise are common problems in biological datasets. Heterogeneity can derive from mixtures of different cell types, such as in sputum, or from sparsity, such as in microbiome or single-cell RNAseq data. Unsupervised methods of dimensionality reduction can effectively eliminate these issues, but suffer from decreased interpretability. That is, variables are collapsed together for reasons that are often opaque. Supervised dimensionality reduction maintains interpretability because variables are collapsed using prior knowledge, such as the genes in a pathway or the expression patterns of a cell type. Here, we combined unsupervised and supervised approaches to de-noise the data while retaining interpretability.

The field is increasingly appreciating the role of the airway microbiome in the development of disease. Commensal microbiota have been shown in other contexts to be strong regulators of host immune system development and homeostasis [26]. Disturbances in the composition of commensal bacteria can result in imbalanced immune responses and affect an individual’s susceptibility to various diseases, including those that are inflammatory (e.g., inflammatory bowel disease and colon cancer), autoimmune (e.g., celiac disease and arthritis), allergic (e.g., asthma and atopy), and metabolic (e.g., diabetes, obesity, and metabolic syndrome) (reviewed in [27]). Investigating the microbiota in the lower respiratory tract is a relatively new field in comparison to the extensive work on the intestinal tract. In fact, the lung was excluded from the original Human Microbiome Project because it was not thought to have a stable resident microbiome [11]. A limited number of reports have investigated the changes in the lung microbiota between healthy, non-smoking and smoking individuals as well as in patients suffering from cystic fibrosis, chronic obstructive pulmonary disease, or asthma [2, 28–30]. Despite emerging data on the airway microbiota, little is known about the role of the lung microbiome in modulating pulmonary mucosal immune responses. LDA-link can find relationships between microbes and genes and link them to immune cells and their responses.

The linkages identified here suggest major processes by which lung immune cells respond to microbes. We found that mast cells respond to *Haemophilus* and *Pasteurella* via IL1B and that eosinophils respond to *Candida* via GCSAML. While experimental validation of these linkages is needed, these results represent observations that would be missed by analyses that do not deconvolve RNAseq data into cell fractions, or that analyze only human RNAseq reads. We expect LDA-link to be broadly useful in relating heterogeneous or noisy RNAseq data.

## METHODS

### Sample collection and sequencing

Sputum induction was performed with hypertonic saline, the mucus plugs were dissected away from the saliva, the cellular fraction was separated, and the RNA was purified as described previously [1]. Briefly, RNA was purified using the All-in-One purification kit (Norgen Biotek) and its integrity was assayed by an Agilent bioanalyzer (Agilent Technologies, Santa Clara, CA). Ribosomal depletion was performed with the RiboGone-Mammalian kit (Clontech Cat. Nos. 634846 & 634847) and cDNA was created with the SMARTer Stranded RNAseq Kit (Cat. Nos. 634836). Samples were sequenced using an Illumina HiSeq 4000 with 2Ô125 bp reads, with an average of 47.5 million reads per sample.

### RNAseq processing by exceRpt

An adapted version of the software package exceRpt [7] was used to process the sputum RNAseq data. Briefly, RNAseq reads were subjected to quality assessment using FastQC software v.0.10.1 (https://www.bioinformatics.babraham.ac.uk/projects/fastqc/) both prior to and following 3’ adapter clipping. Adapters were removed using FastX v.0.0.13 (http://hannonlab.cshl.edu/fastx_toolkit/). Identical reads were counted and collapsed to a single entry and reads containing N’s were removed. Clipped, collapsed reads were mapped directly to the human reference genome (hg19) and pre-miRNA sequences using STAR [31]. Reads that did not align were mapped against a ribosomal reference library of bacteria, fungi, and archaea, compiled by the Ribosome Database Project [32], and then to genomes of bacteria, fungi, plants, and viruses, retreived from GenBank [32]. In cases where RNAseq reads aligned equally well to more than one microbe, a “last common ancestor” approach was used, and the read was assigned to the next node up the phylogenetic tree, as performed by similar algorithms [7, 33].

### Data tables notation

We use the following notation to define matrices associated with *p* patients (115) (Figure 1):

***C***: Clinical table (*N_p_ × N_c_*), c is the clinical index

***G:*** Gene table (*N_p_ × N_g_*), bulk-RNA seq table before deconvolution,

***M***: Microbe abundance table (*N_p_ × N_c_*)

***F***: Cell fractions table (*N_p_ × N_f_*), resulting from the deconvolution of ***G**_p,g_*

***S:*** Cell signatures table (*N_f_ × N_g_*), resulting from the deconvolution of ***G**_p,g_*

***θ**^G^*: Patient topic table (*N_p_ × N_k_*) after LDA inference based on gene table *G_p,g_*

***φ^G^***: Gene topic table (*N_k_ × N_g_*) after LDA inference based on gene table ***G**_p,g_*

***θ**^M^*: Patient topic table (*N_p_ × N_k_*) after LDA inference based on microbe table ***M**_pm_*

***φ**^M^*: Microbe topic table (*N_k_ × N_m_*) after LDA inference based on table ***M**_p,m_*

***L***: gene microbe linkage table (*N_g_ × N_m_*) predicted by LDA-link

### Dimensionality Reduction

#### Supervised, deconvolution

The gene table (**G**) was deconvolved using the transcriptomes from 22 flow cytometry-sorted and sequenced immune cell types (lm22) using the CIBERSORT tool [5]. Briefly, a pre-defined set of characteristic gene expression patterns for each cell type was used to identify the fraction of each cell type given a mixture of expression by solving for the equation:

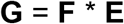

Where **G** is the Gene table of human protein-coding gene expression from the exceRpt pipeline, **F** is the Cell Fraction table, and **E** is the characteristic gene expression calculated within CIBERSORT. Support Vector Regression was used to perform variable selection, reducing the number of characteristic genes used to distinguish cell types and thereby reducing overfitting. The above equation was then solved to provide an estimate of **F**. P-values for the fit of **E** and **F** to **G** demonstrated that all samples were significant at α = 0.05.

Following the solution of **F**, a Cell Signature table **S** was calculated to estimate the expression of *g* genes, as opposed to the reduced set appropriate for the characteristic expression evaluation, by solving the equation:

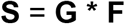

#### Decomposition though a generative model

The Gene table **G** was decomposed using LDA and Non-negative Matrix Factorization (NMF). For LDA, the abundance values for bulk RNAseq and exogenous RNA were scaled down to reduce computation intensity during sampling. More simply, the RPM expression values were converted to integers, and then divided by 10. The max value was set to 1,000.

Given each patient (*p*), all of the genes and microbes were treated like corpus of words in the traditional LDA application. The word (*w*) was gene or microbe, and the word count was gene expression or microbe abundances. We built LDA models for genes and microbes, respectively.

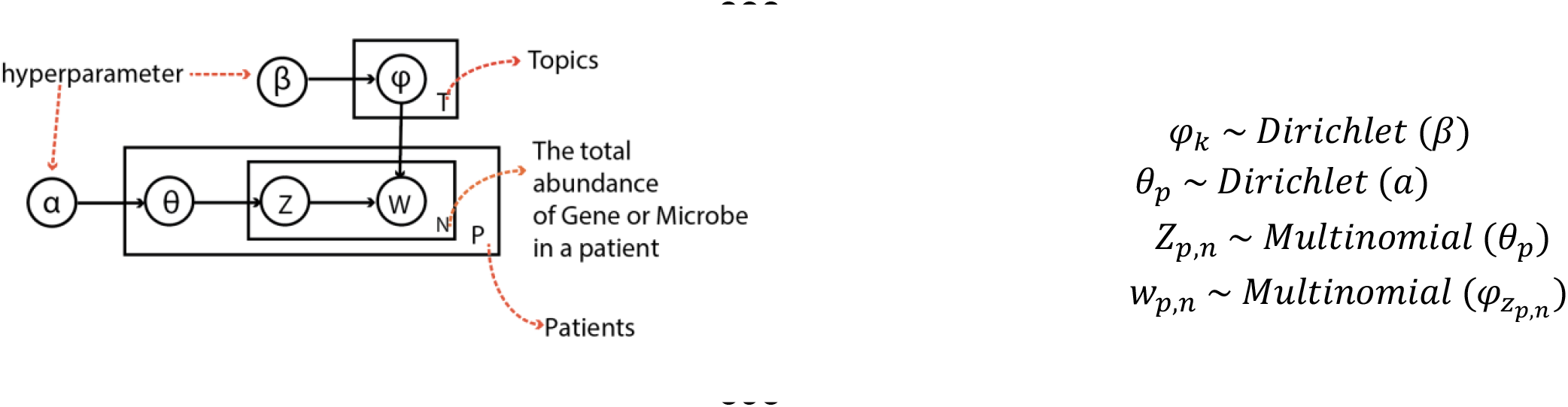

Given *p,w,k,v,N_p_,N_w_,N_k_,N_v_*, α, β, Z, θ, φ,W, where *p,w,k,v* denote a patient, a word in a document, a topic and a word in the corpus respectively; *N_p_* is the number of documents(patients), N_*w*_ is the number of words (gene or microbe) in *a* document, N_*k*_ is the number of topics (set as 10), N_*v*_ is the corpus for all the documents; α (*N_k_* dimensional vector) and β (*N_v_*-dimension al vector) are the hyper parameters for θ (*N_p_* × *N_k_*, the distributoin of topics in documents) and φ (*N_k_ × N_v_*, the distribution of word for topics) W is an *N_w_*-dimensional vector that denotes the word (gene or microbe expression) in a document (patients). Z is the *N_w_*-dimensional vector of integers between 1 and *N_k_* for the topic of word in a document.

The joint distribution of the LDA model is 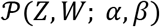 and φ and θ are integrated out as:

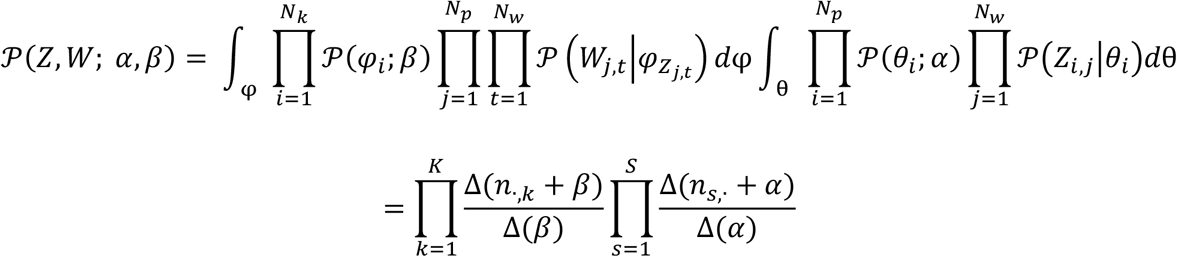

Where 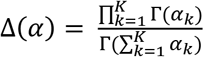

Gibbs sampling equation can be derived from 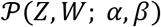 to approximate the distribution of 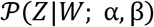 because 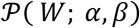 is invariant to *Z*. Given *Z_m,n_* denotes the topic of the *n*th word token in the *m*th document, and also assume that its word symbol is the *v*th word in the vocabulary, the conditional probability can be inferred as follows:

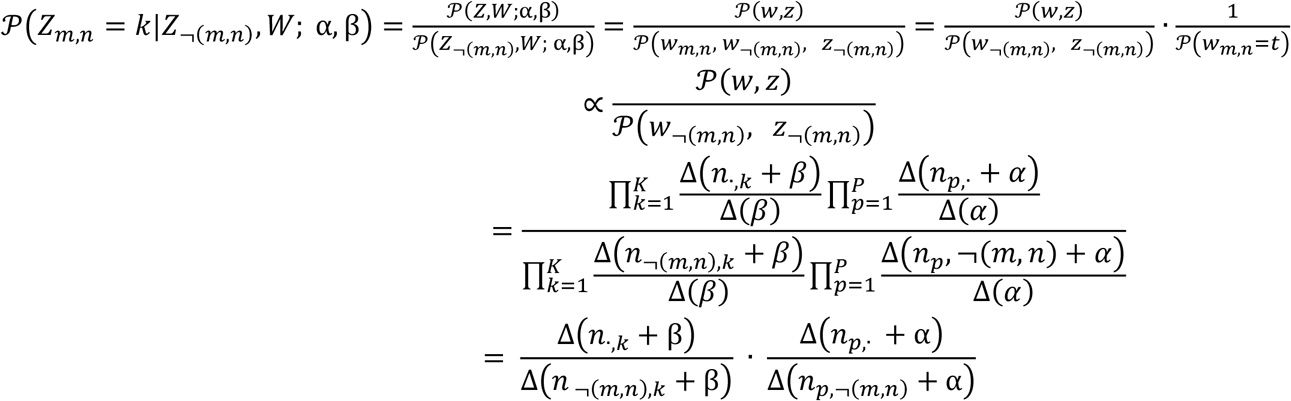

After sampling, the expectation of the θ (doc → topic) and φ(topic → word) matrix can be inferred as follows given the symmetric hyper-parameters *α* and *β* were used:

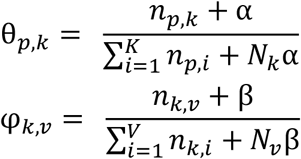

We instantiated the variables θ and φ to 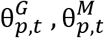, and 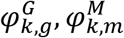, where 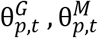 denotes the gene and microbe topic fraction in patient; 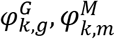 denotes the gene and microbe topic.

### Single-cell RNAseq

Sputum cells were separated on a Fluidigm C1 medium-sized channel. The mRNA was purified from approximately 500pg-1ng of total RNA using the Clontech SMARTer Ultra Low RNA Kit and poly-dA-selected using SPRI beads and dT primers. Full-length cDNA was sheared into 200-500bp DNA fragments by sonication (Covaris, Massachusetts, USA), and then indexed and size validated by LabChip GX. Two nM libraries were loaded onto Illumina version 3 flow cells and sequenced using 75bp single-end sequencing on an Illumina HiSeq 2000 according to Illumina protocols. Data were cleaned, processed, aligned, and quantified following the SINCERA pipeline [34].

### Pathogen-to-host linkage identification

Microbe relative abundances and gene TPM values were correlated as follows, with *G·,_i_* for *i* gene and *M·,_j_* for *j* microbe:

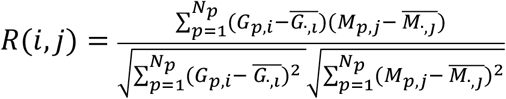

Gene-microbe correlations with *p*-values less than 1e-5 (absolute correlation greater than 0.4) were chosen as the positive links in a training set. Negative links in the training set were defined as an absolute correlation of less than 0.05. This approach resulted in 302 positive and 650,398 negative links. A random forest algorithm was trained on this set, which can accommodate the highly unbalanced dataset as well as potentially identify non-linear links between genes and microbes. Down-sampling and up-sampling techniques were tested but did not significantly improve the model. In the final model, we adopted the upscaling technique and tested it using cross-validation. The positive dataset was upscaled to very high levels. We use 2-fold cross validation to validate the performance. Simply, we randomly select half training data to train the model, and use the remaining records to test the performance and repeat this for ten times. The AUC and AUPR were 0.994 and 0. 996 on average, respectively.

### Microbe co-abundance network

The raw abundance *M* and LDA microbe topic matrices *φ^M^*, which represent the microbe’s weight to each topic, were generated.

The correlation network between different microbes was calculated using Pearson correlation. The cutoff to define a co-abundance edge was 0.8 for 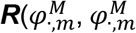 and 0.3 for ***R***(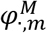. The microbe network modules, which were densely connected themselves but sparsely connected to other modules, were clustered based on between-ness [35] and the other algorithms; we also tested label propagation and fast greedy algorithms [36].

We also compared the LDA topics with the microbes in the same clusters. If a microbe was the top 10 most highly contributed for a topic, then we labeled the topic number in the bracket. Some microbes may have multiple topic labels because they highly contribute to multiple topics.

## DATA ACCESS

Sputum bulk-cell RNAseq data can be found under the bioproject SRRXXXXX and sputum single-cell RNAseq data at SRRYYYYYY.

## ACKNOWLEDGMENTS

This work was supported by a National Library of Medicine fellowship to DS (5T15LM007056-28) and an NHLBI grant to GC (1R01HL118346-01). The authors would like to thank the support of the Yale High Performance Computing services (Grace, Ruddle, Farnam) and Yale Center for Genome Analysis.

## ABBREVIATIONS

LDA: Latent Dirichlet Allocation
PCA: Principle Component Analysis

## SUPPLEMENTAL INFORMATION

**Figure S1:**

The distribution of cell fraction of sample in different group.

**Figure S2**

The heatmap between NMF component and cell fraction from LM22.

**Figure S3**

Overall view of correlation of all the extracellular organism with clinical features.

**Figure S4**

Heatmap of the correlation of topics (gene and microbe) with clinical information.

**Figure S5**

Co-abundance network based on correlation of abundance.\

**Figure S6**

Top associated microbes in microbe topics

Figure S7

Top associated genes in gene topics

Figure S8

(A) The topic distribution of patient. (B) The gene enrichment analysis of top genes in topic 4.

Figure S9

Main pathways get involved by microbe linked genes.

