## Supplemental Figures for "Approaches for integrating heterogeneous RNA-seq data reveals cross-talk between microbes and genes in asthmatic patients"

Supplementary information

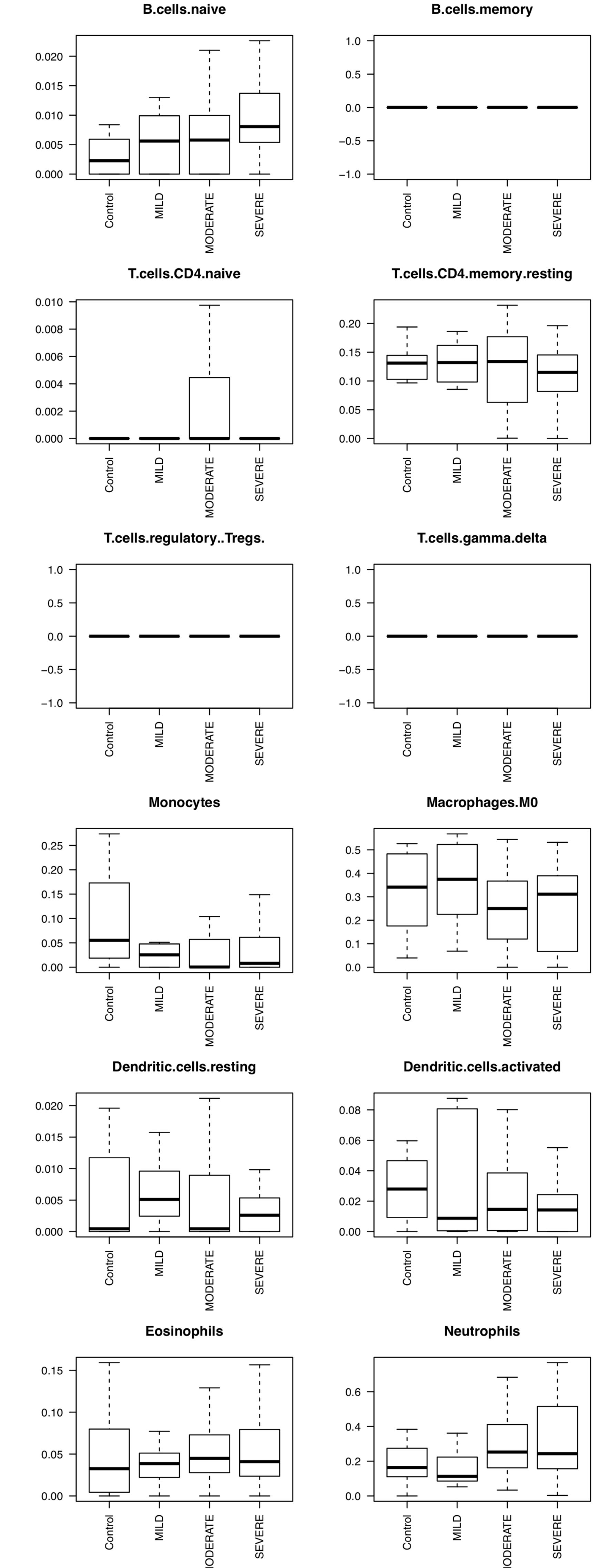


### Figure S1

Boxplot for cell-type frequency in four patient groups: Control, mild, moderate and severe

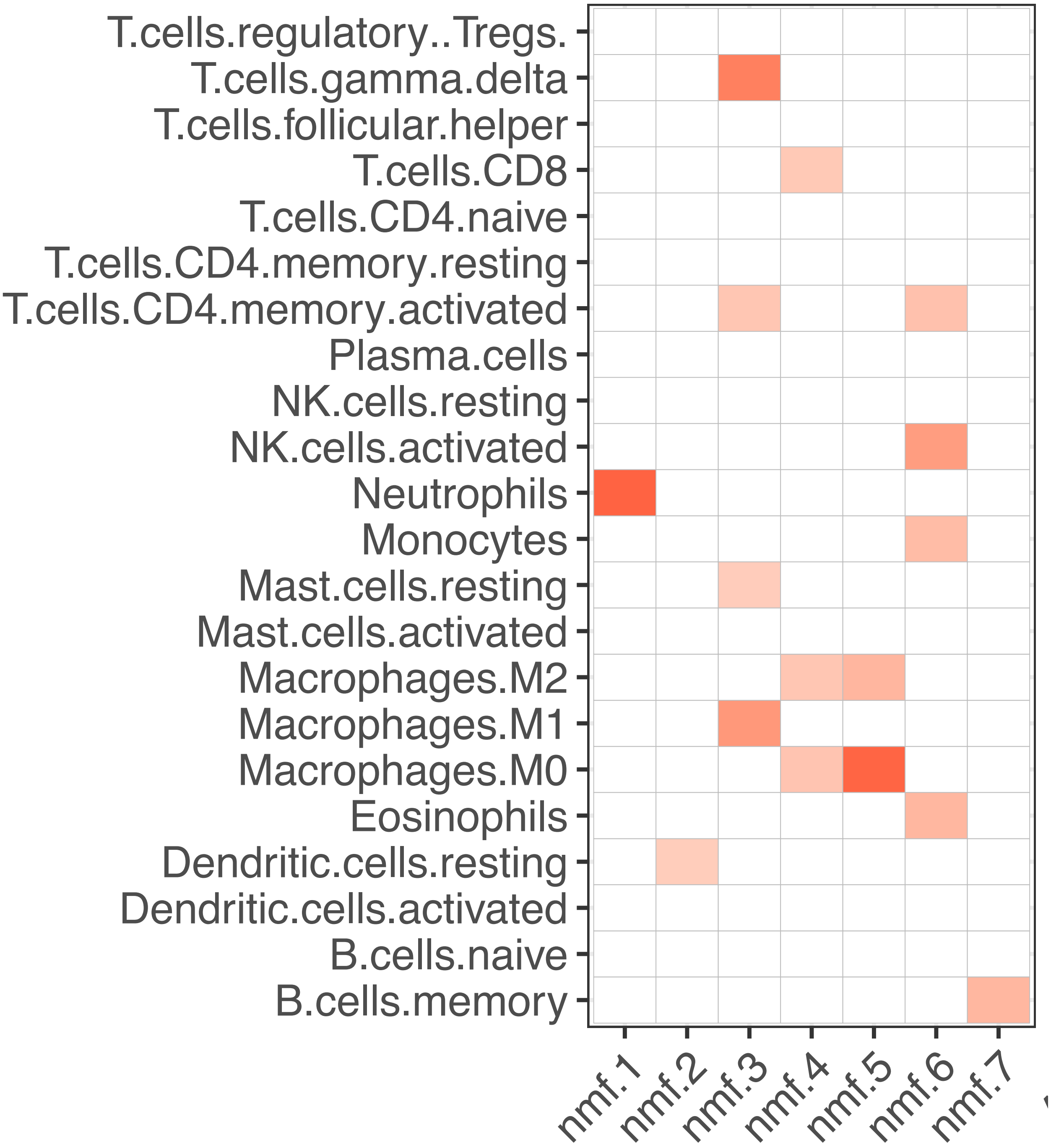


Figure S2:

Correlation between NMF weight matric and cell-type fraction

Figure S3: microbe abundance correlation with clinical information.

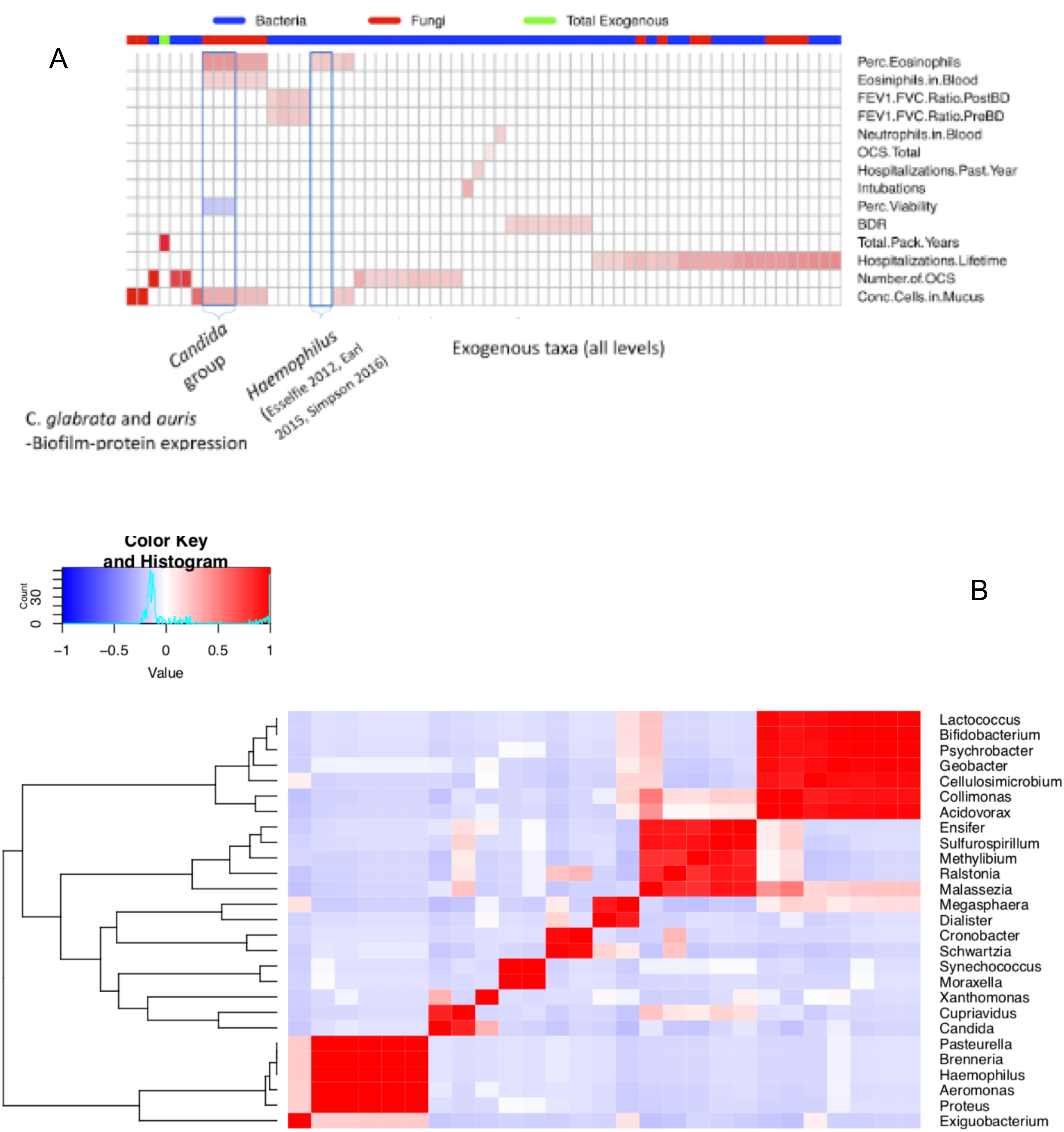

Figure S4: correlation between gene and microbe topic fraction in patients and clinical information.

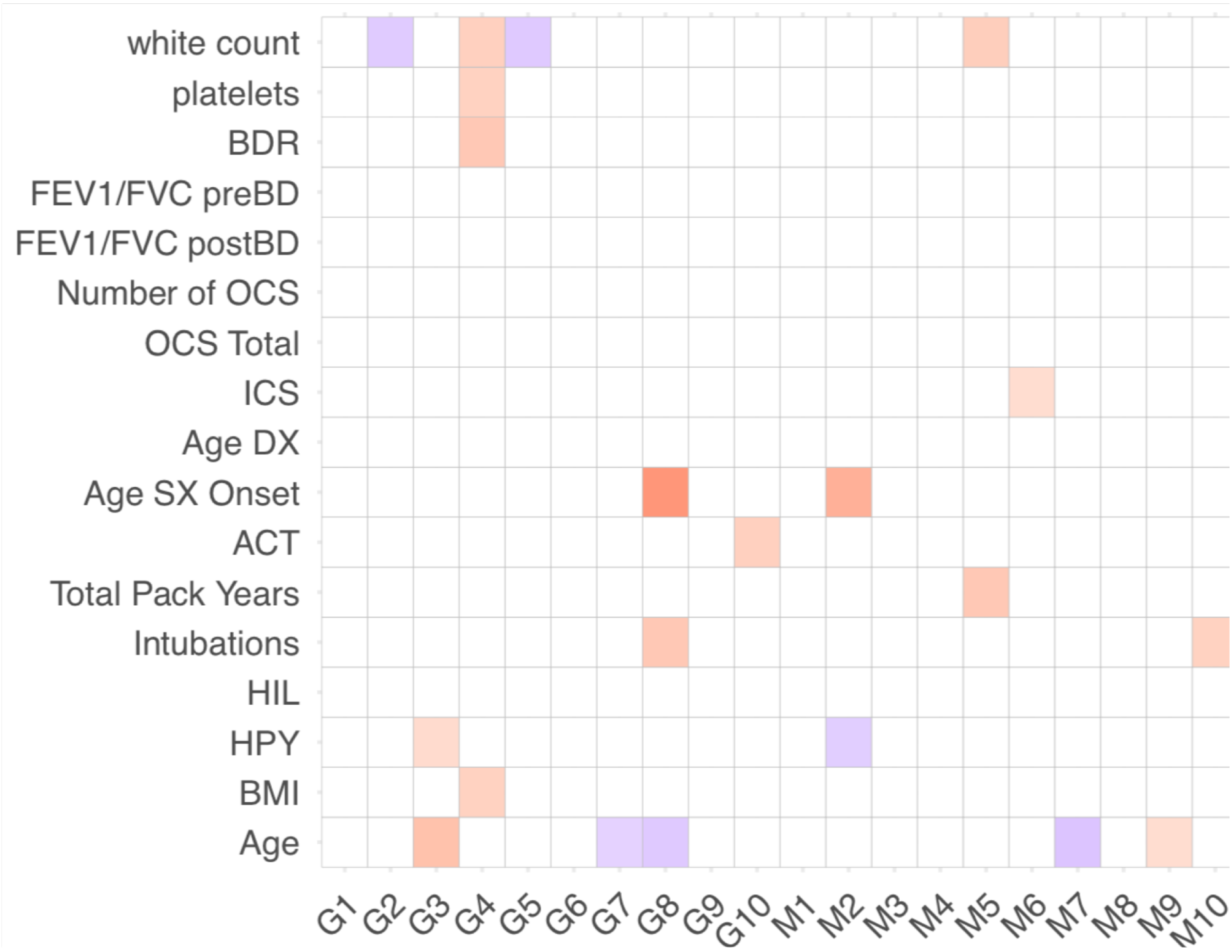

Figure S5: microbe co-abundance network using the correlation with raw microbe abundance (A: in network view; B: heatmap view).

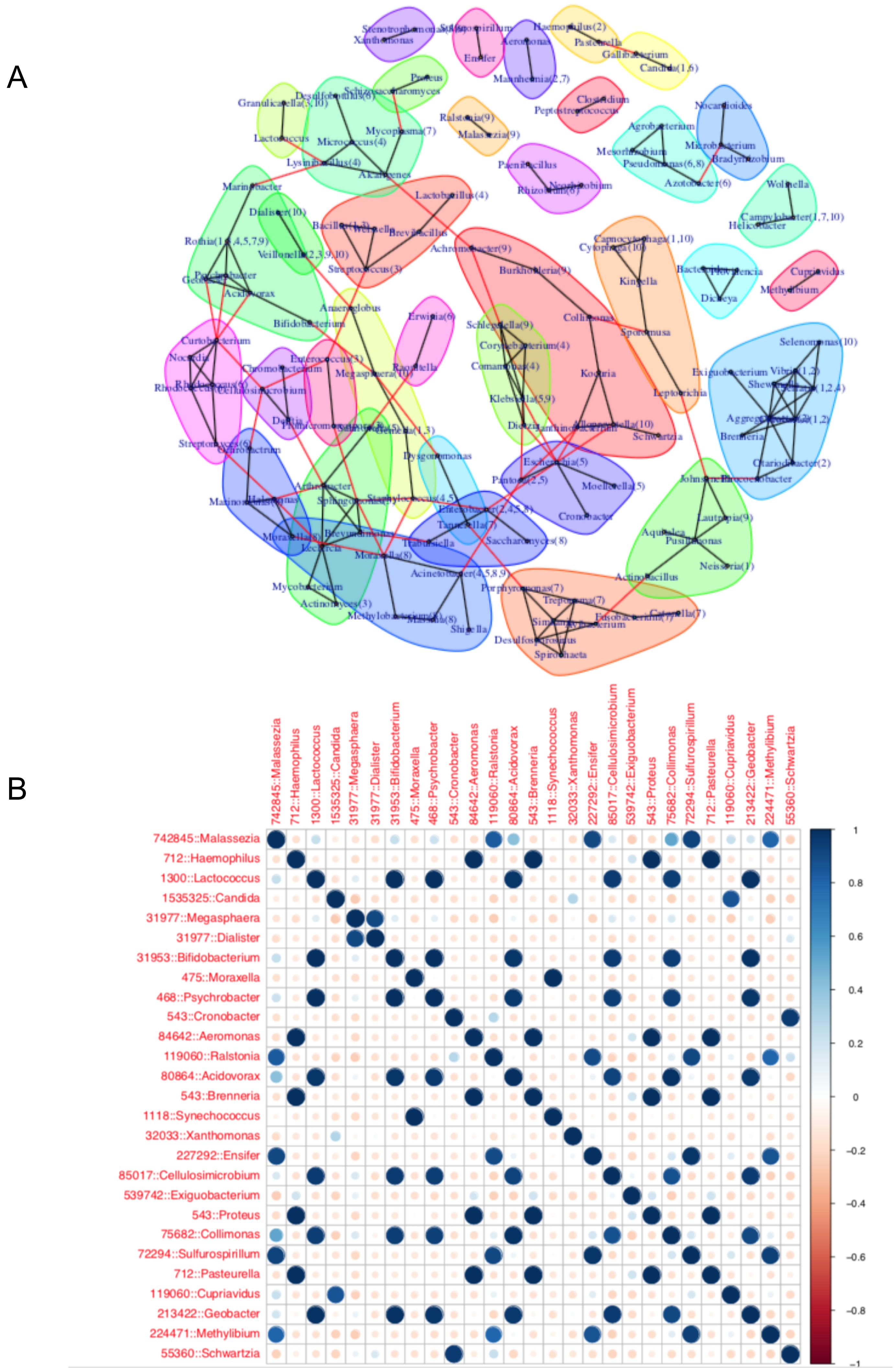

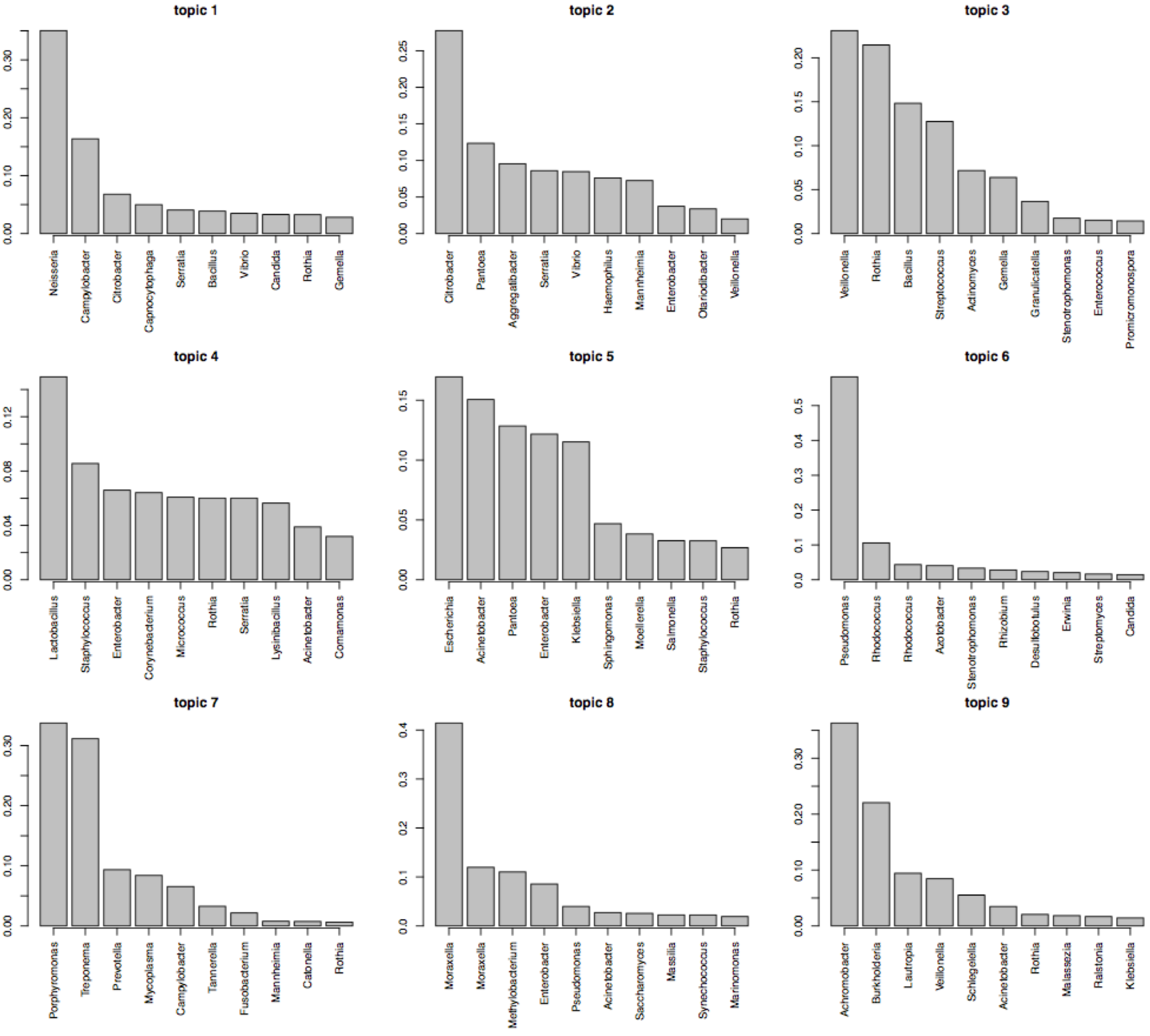


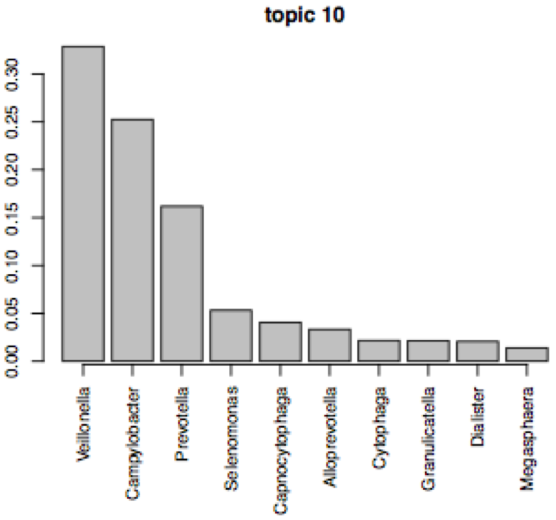


Figure S6
Top weighted microbe in the microbe topics.

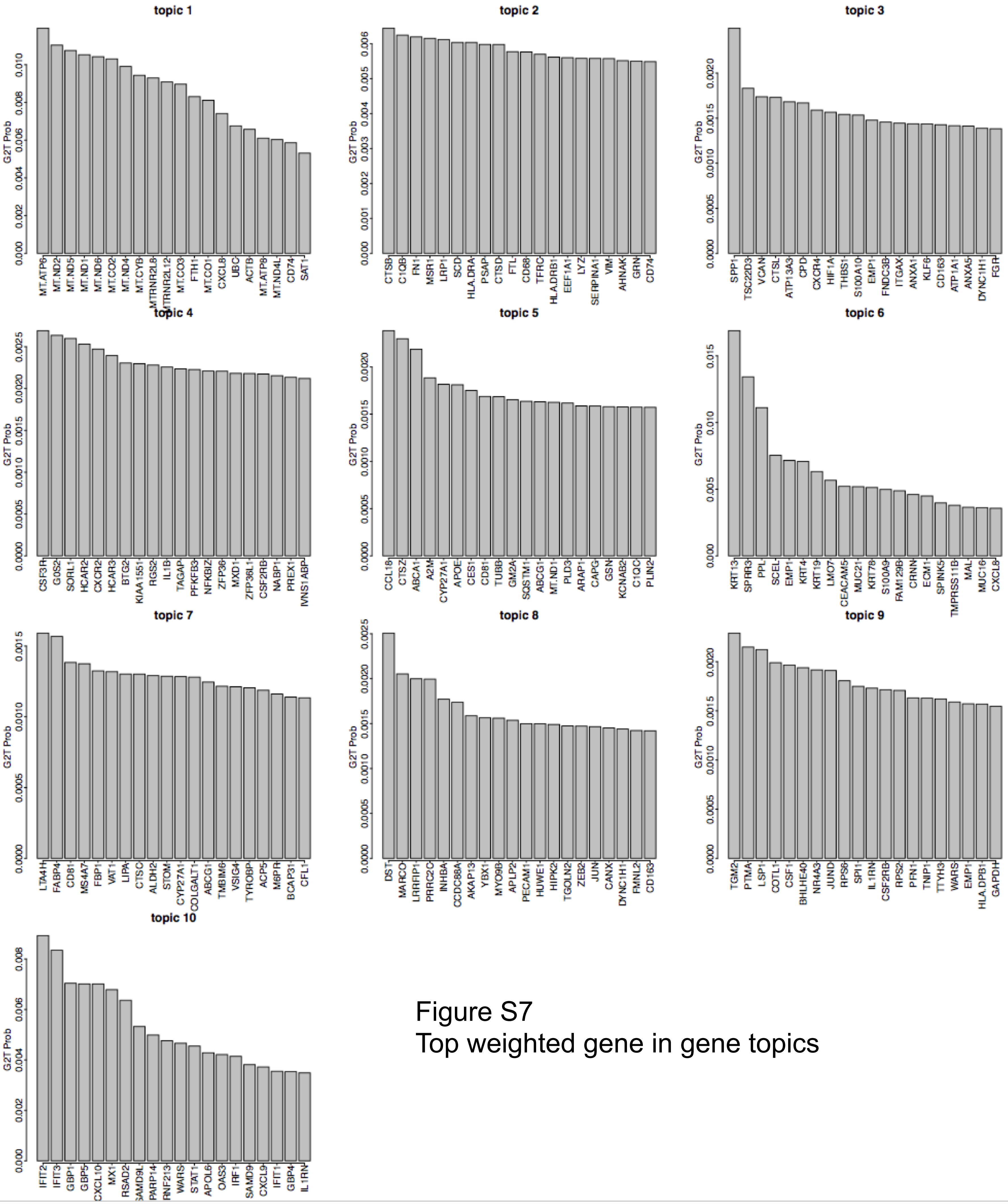


Figure S7
Top weighted gene in gene topics

Figure S8:
Heatmap of Gene topic fraction in patient (A) and
Gene enrichment analysis of top weighted gene in topic 4 (B).

A

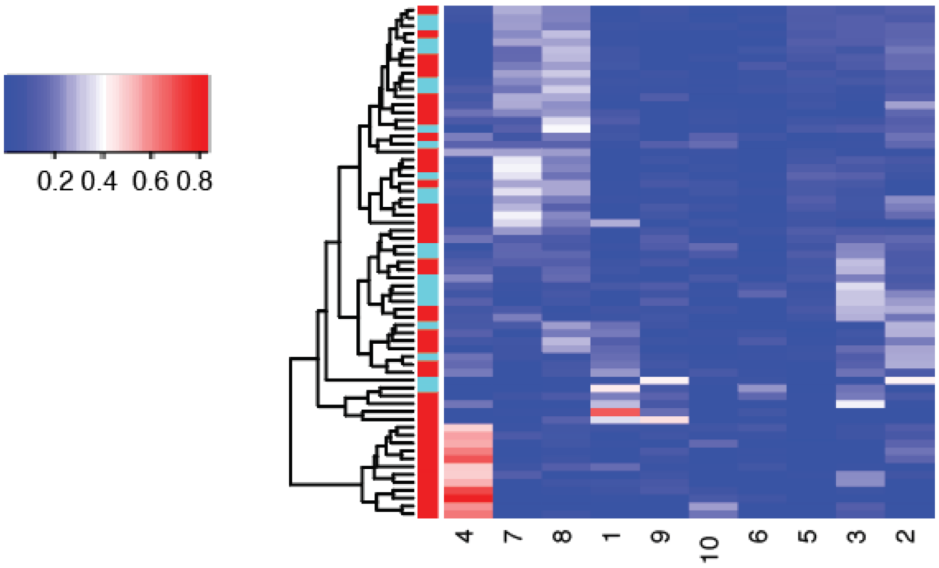


B

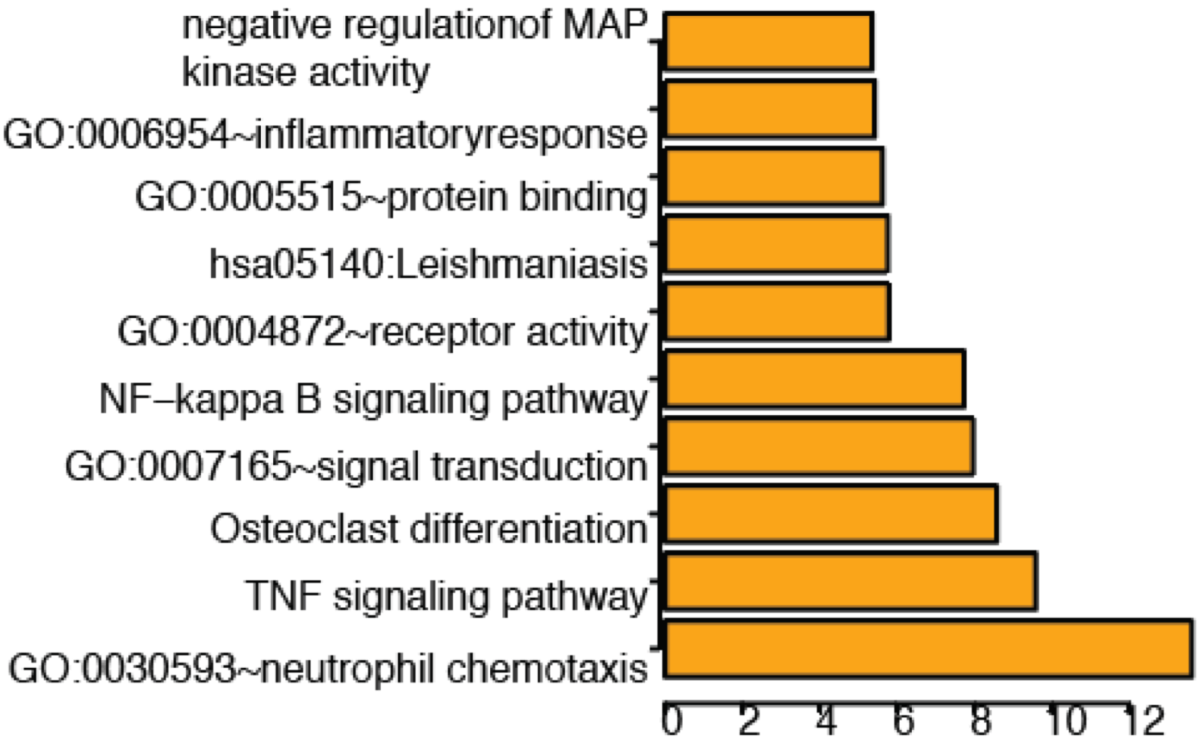

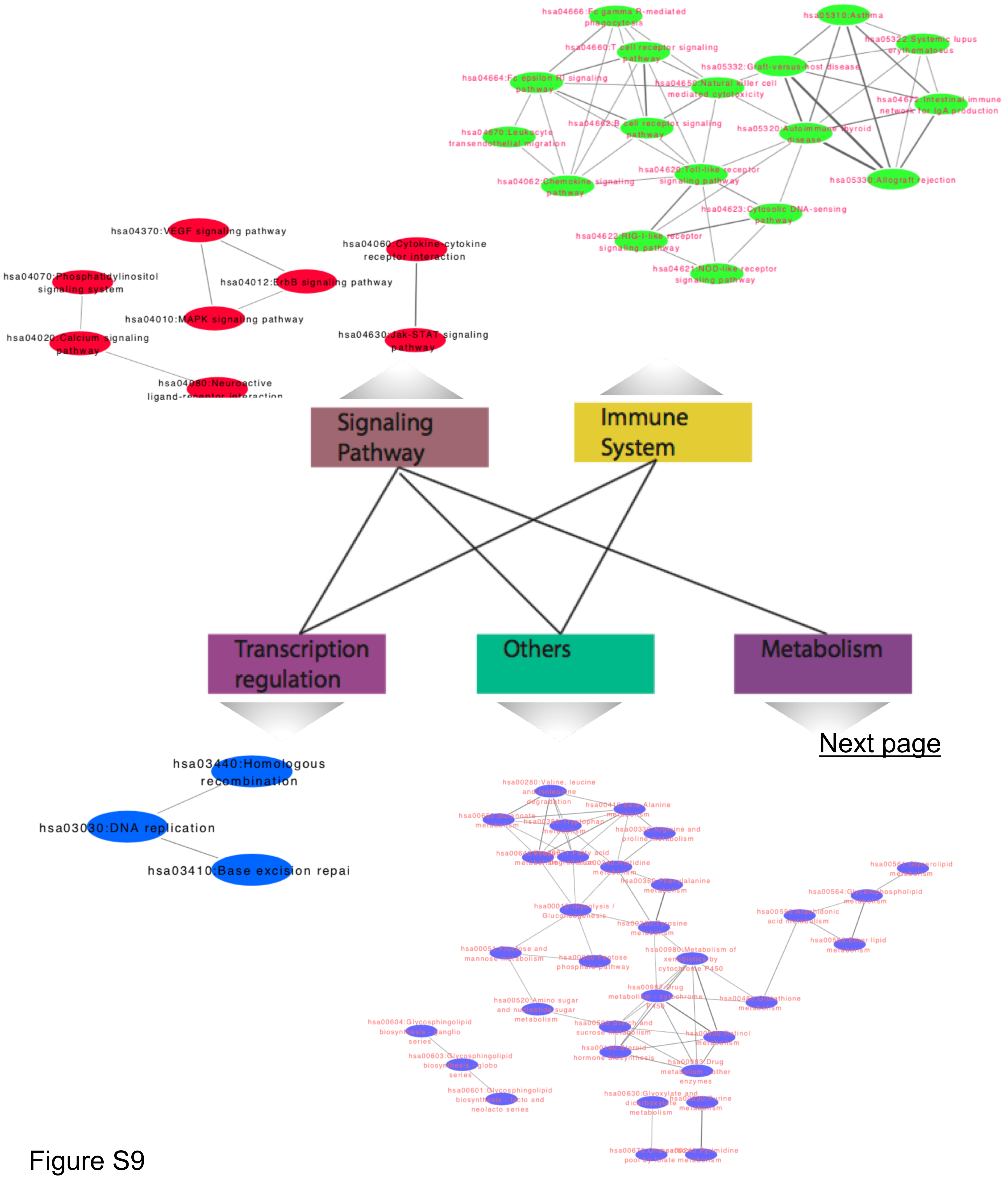


Figure S9

Function annotation finds the enriched gene pathway

Figure S9 continued: Metabolism network.

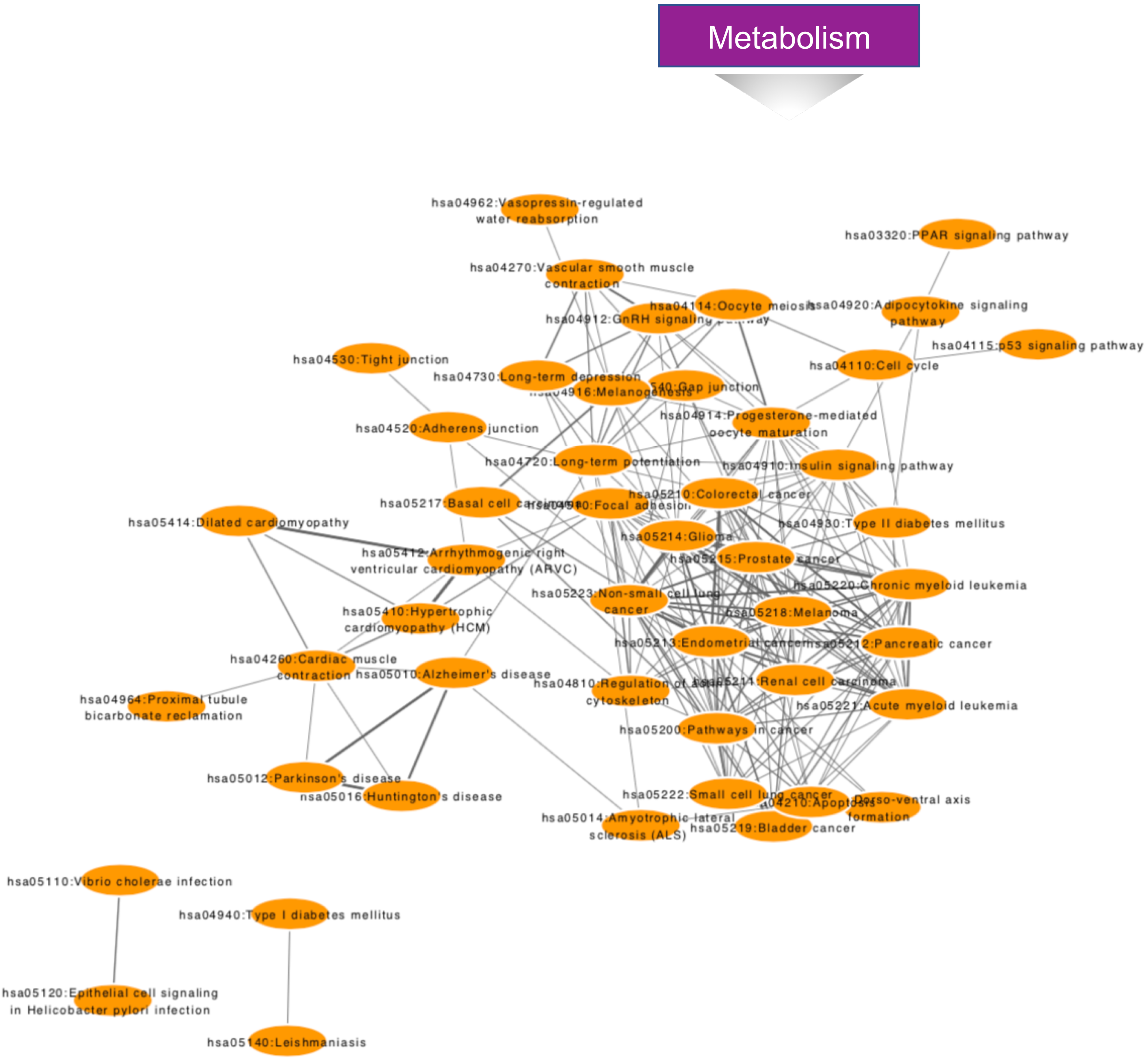
